# Sensitivity and uncertainty analysis of two human atrial cardiac cell models using Gaussian process emulators

**DOI:** 10.1101/818047

**Authors:** Sam Coveney, Richard H. Clayton

## Abstract

Cardiac cell models reconstruct the action potential and calcium dynamics of cardiac myocytes, and are becoming widely used research tools. These models are highly detailed, with many parameters in the equations that describe current flow through ion channels, pumps, and exchangers in the cell membrane, and so it is difficult to link changes in model inputs to model behaviours. The aim of the present study was to undertake sensitivity and uncertainty analysis of two models of the human atrial action potential. We used Gaussian processes to emulate the way that 11 features of the action potential and calcium transient produced by each model depended on a set of. The emulators were trained by maximising likelihood conditional on a set of design data, obtained from 300 model evaluations. For each model evaluation, the set of inputs was obtained from uniform distributions centred on the default values for each parameter, using latin-hypercube sampling. First order and total effect sensitivity indices were calculated for each combination of input and output. First order indices were well correlated with the square root of sensitivity indices obtained by partial least squares regression of the design data. The sensitivity indices highlighted a difference in the balance of inward and outward currents during the plateau phase of the action potential in each model, with the consequence that changes to one parameter can have opposite effects in the two models. Overall the interactions among inputs were not as important as the first order effects, indicating that model parameters tend to have independent effects on the model outputs. This study has shown that Gaussian process emulators are an effective tool for sensitivity and uncertainty analysis of cardiac cell models.

**Author summary:** The time course of the cardiac action potential is determined by the balance of inward and outward currents across the cell membrane, and these in turn depend on dynamic behaviour of ion channels, pumps and exchangers in the cell membrane. Cardiac cell models reconstruct the action potential by representing transmembrane current as a set of stiff and nonlinear ordinary differential equations. These models capture biophysical detail, but are complex and have large numbers of parameters, so cause and effect relationships are difficult to identify. In recent years there has been an increasing interest in uncertainty and variability in computational models, and a number of tools have been developed. In this study we have used one of these tools, Gaussian process emulators, to compare and contrast two models of the human atrial action potential. We obtained sensitivity indices based on the proportion of variance in a model output that is accounted for by variance in each of the model parameters. These sensitivity indices highlighted the model parameters that had the most influence on the model outputs, and provided a means to make a quantitative comparison between the models.

## Introduction

The cardiac action potential arises from the movement of ions through channels, pumps, and exchangers in the cell membrane. At any instant, the current carried by each ionic species depends on potential difference across the cell membrane, as well as ion concentrations and the dynamics of ion channel gating. The complex interplay of currents produces depolarisation and then repolarisation of the membrane, which then acts to trigger release of Ca^2+^, initiating mechanical contraction.

The first model of the action potential in a cardiac myocyte was developed over 50 years ago [1], and since then a series of more detailed models have been developed as experimental techniques and data have improved. The present generation of models provide detailed reconstructions of the cardiac action potential [2], and computational models of cardiac cells and tissue have become valuable research tools because they can encode biophysical mechanisms into a quantitative framework, and so can be used to test and construct hypotheses [3].

Although these detailed models are capable of simulating the behaviour of real cardiac myocytes, this veracity comes at the price of complexity. Models of the cardiac action potential typically comprise a system of coupled, stiff, and non-linear ordinary differential equations. There are many model parameters and boundary conditions, which we will refer to as *model inputs* from here onward. These model inputs can be derived from experimental data, using approaches based on those pioneered by Hodgkin and Huxley in squid giant axon [4]. However, experimental data are subject to variability and error arising from both limitations of experimental methods as well as intrinsic variability in cardiac cells. Some of these inputs, such as binding affinities and reaction rate constants, can be considered to have fixed values because they have a physical basis. However others, such as maximum ion channel conductance, depend on the ion channel density in the cell membrane as well as other factors that are variable. These quantities may therefore vary from one cell to another, and even from beat to beat in the same cell. These considerations underlie three specific problems. First, errors and variability in data are typically not taken into account when fitting model inputs, and taking an average of experimental data can distort model behaviour [5]. Second, data from different sets of experiments can result in different models of the same cell type. For example, there are several models of the human atrial action potential, all based on human data, but which show different types of behaviour [6, 7]. Finally, a further complication arises from the modular nature of cardiac cell models. The equations for a particular ion channel, pump, or exchanger are often re-used in different models and so the provenance of model inputs may be very difficult to establish [8].

Addressing these problems requires approaches that can establish how model behaviours and outputs depend on inputs that may be uncertain. However, the level of detail included in the present generation of cardiac cell models means that formal analysis is difficult, and so a detailed examination of how model inputs influence outputs relies on large numbers of numerical simulations where the inputs vary from one model run to the next [9]. These datasets can be used for regression analysis, which enables the sensitivity of model outputs to changes in model inputs to be assessed [10]. Another approach is to select a set of inputs, or population of models, that produce action potentials that are in the range of experimental observations [11–13]. A drawback of these approaches arises from the high dimensional input space for the models; a very large number of model evaluations is needed to investigate the input space thoroughly [14], although recent work indicates that this challenge can be overcome by new models that are designed for uncertainty quantification [15].

Methods to quantify uncertainty and variability using probabilistic approaches have been developed and applied in a number of areas [16], including cardiovascular flow models [17]. These approaches are promising because an emulator of a computational model can treat uncertainties explicitly, and can be evaluated at much lower computational cost than the original model [19, 42]. Gaussian Processes are flexible non-parametric regression tools capable of fitting complex data, thus they are widely used in the machine learning community. They provide confidence intervals on their predictions, and are therefore ideal tools for sensitivity and uncertainty analysis.

In this report we demonstrate the use of Gaussian process emulators for sensitivity and uncertainty analysis of two biophysically detailed models of the human atrial action potential. Our objectives were to gain insight into the comparative mechanisms of each model by first calculating variance based sensitivity indices that quantify how variance in model inputs influences variance in model outputs, and second by examining how each model behaved under uncertain inputs.

## Materials and methods

### Human atrial cell models

Several models of the human atrial action potential have been developed and are reviewed in detail elsewhere [6, 7, 12]. We selected two models for the present study, both based on data from human atrial cells. The first model is the *Courtemanche* model [20]. The second model is an extension of the model developed by Nygren et al [21], with modifications to the *I*_*Kur*_ and *I*_*to*_ currents as well as the movement of *Na*^+^ [22], which we refer to as the *Maleckar* model.

We chose these models because both represent the action potential of human atrial cells, and so have clinical relevance. Both have been used for tissue and whole-organ scale simulations of atrial fibrillation [23–26]. Furthermore, both models have a comparable set of ion channels, pumps, and exchangers, however differences in *Ca*^2+^ handling underlie the different action potential shapes [7].

### Model inputs and outputs

The Courtemanche and Maleckar cell models include components that represent membrane electrophysiology as well as intracellular *Ca*^2+^ storage, uptake, and release. We chose to concentrate on parameters that control the maximum current density carried by ion channels, pumps, and exchangers in the cell membrane as well as those that control the rate and magnitude of uptake and release of intracellular *Ca*^2+^. We also selected the cell capacitance *C*_*m*_, and the extracellular concentrations [*Na*^+^]_*o*_, [*K*^+^]_*o*_, and [*Ca*^2+^]_*o*_ as additional inputs. The inputs examined in this study are listed in Table 1, where the central values given are the default for each model.

**Table 1.** Range of inputs used for design data in each cell model.

| **Courtemanche model** |  |  |  |
| --- | --- | --- | --- |
| Input | Central value | Range | Units |
| $G_{Na}$ | 7.8 | 5.85 - 11.70 ($\pm 50\%$) | nS/pF |
| $G_{K1}$ | 0.09 | 0.045 - 0.018 ($\pm 25\%$) | nS/pF |
| $G_{to}$ | 0.165 | 0.0826 - 0.0330 ($\pm 50\%$) | nS/pF |
| $fG_{Kur}$ | 1.0 | 0.50 - 1.50 ($\pm 50\%$) | none |
| $G_{Kr}$ | 0.0294 | 0.0147 - 0.0441 ($\pm 50\%$) | nS/pF |
| $G_{Ks}$ | 0.1294 | 0.0647 - 0.1941 ($\pm 50\%$) | nS/pF |
| $G_{Ca,L}$ | 0.1237 | 0.0619 - 0.1856 ($\pm 50\%$) | nS/pF |
| $G_{b,Na}$ | 0.0006 | 0.0003 - 0.0010 ($\pm 50\%$) | nS/pF |
| $G_{b,Ca}$ | 0.0011 | 0.0005 - 0.0017 ($\pm 50\%$) | nS/pF |
| $i_{NaKMax}$ | 0.5993 | 0.2997 - 0.8990 ($\pm 50\%$) | pA/pF |
| $i_{NaCaMax}$ | 1600.0 | 800.00- 2400.0 ($\pm 50\%$) | pA/pF |
| $i_{p,CaMax}$ | 0.275 | 0.1375 - 0.4125 ($\pm 50\%$) | pA/pF |
| $K_{rel}$ | 0.30 | 0.15 - 0.45 ($\pm 50\%$) | /ms |
| $\tau_{tr}$ | 180.0 | 90.0 - 270.0 ($\pm 50\%$) | ms |
| $i_{upMax}$ | 0.005 | 0.0025 - 0.0075 ($\pm 50\%$) | mM/ms |
| $K_{up}$ | 0.00092 | 0.00046 - 0.0014 ($\pm 50\%$) | mM |
| $C_m$ | 100.0 | 75.0 - 125.0 ($\pm 25\%$) | pF |
| $[Na^+]_o$ | 140.0 | 126.0 - 154.0 ($\pm 10\%$) | mM |
| $[K^+]_o$ | 5.4 | 4.86 - 5.94 ($\pm 10\%$) | mM |
| $[Ca^{2+}]_o$ | 1.8 | 1.62 - 1.98 ($\pm 10\%$) | mM |
| **Maleckar model** |  |  |  |
| Input | Central value | Range | Units |
| $P_{Na}$ | 0.0018 | 0.0009 - 0.0027 ($\pm 50\%$) | nL/s |
| $G_{K1}$ | 3.1 | 2.325 - 3.875 ($\pm 25\%$) | nS |
| $G_t$ | 8.25 | 4.125 - 12.375 ($\pm 50\%$) | nS |
| $G_{Kur}$ | 2.25 | 1.125 - 3.375 ($\pm 50\%$) | nS |
| $G_{Kr}$ | 0.5 | 0.250 - 0.750 ($\pm 50\%$) | nS |
| $G_{Ks}$ | 1.0 | 0.50 - 1.50 ($\pm 50\%$) | nS |
| $G_{Ca,L}$ | 6.75 | 3.375 - 10.125 ($\pm 50\%$) | nS |
| $G_{b,Na}$ | 0.0605 | 0.303 - 0.909 ($\pm 50\%$) | nS |
| $G_{b,Ca}$ | 0.0590 | 0.0295 - 0.0885 ($\pm 50\%$) | nS |
| $NaKMax$ | 68.55 | 34.27 - 102.82 ($\pm 50\%$) | pA |
| $K_{NaCa}$ | 0.0750 | 0.0375 - 0.1125 ($\pm 50\%$) | pA/mM<sup>4</sup> |
| $i_{p,CaMax}$ | 4.0 | 2.0 - 6.0 ($\pm 50\%$) | pA |
| $\alpha_{rel}$ | 200000 | 100000 - 300000 ($\pm 50\%$) | pA/mM |
| $\tau_{tr}$ | 0.01 | 0.005 - 0.015 ($\pm 50\%$) | s |
| $i_{upMax}$ | 2800 | 1400 - 4300 ($\pm 50\%$) | pA |
| $K_{cyca}$ | 0.0003 | 0.00015 - 0.00045 ($\pm 50\%$) | mM |
| $K_{srca}$ | 0.5 | 0.25 - 0.75 ($\pm 50\%$) | mM |
| $K_{xcs}$ | 0.4 | 0.2 - 0.6 ($\pm 50\%$) | dimensionless |
| $C_m$ | 50 | 37.5 - 62.5 ($\pm 25\%$) | pF |
| $[Na^+]_o$ | 140.0 | 126.0 - 154.0 ($\pm 10\%$) | mM |
| $[K^+]_o$ | 5.4 | 4.86 - 5.94 ($\pm 10\%$) | mM |
| $[Ca^{2+}]_o$ | 1.8 | 1.62 - 1.98 ($\pm 10\%$) | mM |

The rationale for this choice was that each of the selected inputs can be considered uncertain (*i.e.* not a physical constant), yet has a biophysical interpretation. Maximum conductances of ion channels, pumps, and exchangers depend on protein expression, and so could be expected to vary within an individual cell at different times as well as from cell to cell. Cell size and capacitance vary from cell to cell. This natural variability can be considered to be aleatoric uncertainty, which is irreducible [16]. On the other hand, the kinetics of transmembrane currents are related to ion channel biophysics, and so could be considered epistemic uncertainty, which can in principle be reduced.

Our analysis proceeded in two stages. In *Stage 1* the influence of all of the inputs listed in Table 1 was examined for a fixed pacing cycle length of 1000 ms. In *Stage 2*, a subset of the inputs was selected on the basis of their *Stage 1* sensitivity index (see below), and a new set of emulators was built using this subset as inputs, together with the diastolic interval (DI) of the S2 beat in an S1-S2 pacing sequence as an additional input.

Cardiac cell models produce an output that is a time series of states. Of these, membrane voltage *V*_*m*_ and intracellular *Ca*^2+^ concentration [*Ca*^2+^]_*i*_ describe the time course of action potentials and *Ca*^2+^. To investigate how cell model inputs influence action potential shape, we selected nine features of the action potential that quantify its shape, based on biomarkers used in related work [11, 12] as well as the minimum and maximum intracellular *Ca*^2+^ concentration. These eleven outputs are shown in Fig 1 and are listed below.

- *dV*_*m*_/*dt*_*max*_ – Maximum slope of the action potential upstroke.
- *V*_*max*_ – Peak voltage of the action potential.
- *V*_20_, *V*_40_, *V*_60_, and *V*_80_ – Membrane voltage measured at 20, 40, 60, and 80% of *APD*_90_.
- *APD*_50_ and *APD*_90_ – Action potential duration at 50% and 90% of repolarisation.
- *RestV*_*m*_ – Resting membrane potential, calculated as the average membrane voltage over a 10 ms period, 100 ms prior to the action potential upstroke.
- 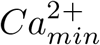 and 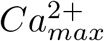 – Minimum and maximum intracellular Ca^2+^ concentration.

**Fig 1.**
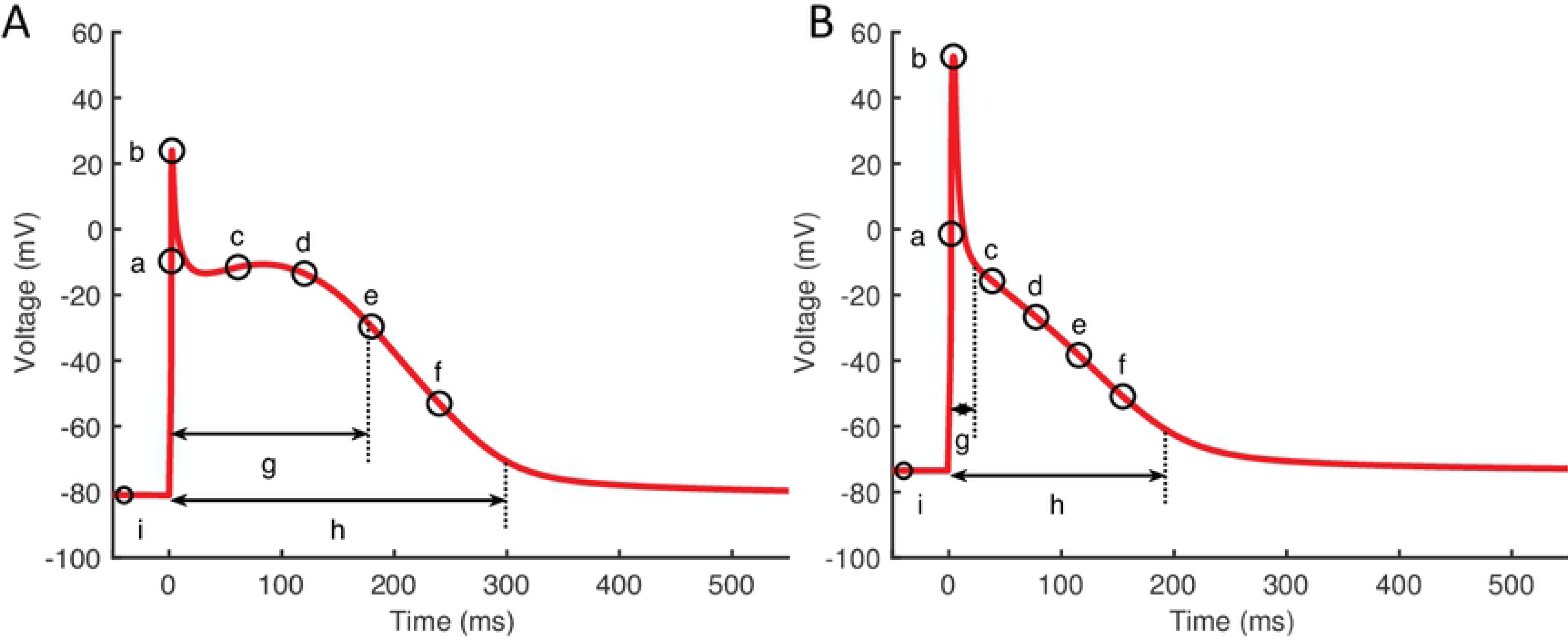
Action potential biomarkers. Nine action potential biomarkers were used as outputs to characterise each simulator run. A: Courtemanche model, B: Maleckar model. Biomarker labels are a: *dV/dt*_*max*_, b: *V*_*max*_, c-f: *V*_20_, *V*_40_, *V*_60_, and *V*_80_, g: *APD*_50_, h: *APD*_90_, i: *RestV*_*m*_. 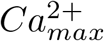 and 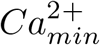 not shown

### Model implementation

Both cell models were implemented in Matlab (Mathworks inc.), using Matlab code automatically generated from the CellML repository (http://cellml.org). The models were solved using the ode15s time adaptive solver for stiff systems of ODEs with the relative and absolute tolerance both set to 10^−6^, and the maximum time step set to 0.5 ms.

To ensure that both cell models remained stable over the range of inputs used to train the GP emulators, we made some small modifications. Previous studies (for example [6]) have identified an instability in the Courtemanche model that arises from a gradual drift in the intracellular concentrations [Na^+^]_i_ and [*K*^+^]_*i*_. We therefore fixed [*Na*^+^]_*i*_ and [*K*^+^]_*i*_ at their default initial values of 11.17 and 139.00 mM respectively in the Courtemanche model implementation. In the Maleckar model, we fixed the *I*_*K,Ach*_ current to zero.

For each run, action potentials were initiated by an inward current of 2000 pA delivered for 2 ms in the Courtemanche model, and 750/C_m_ pA/pF for 6 ms in the Maleckar model. In *Stage 1*, each run comprised 40 action potentials at a cycle length of 1000 ms. In *Stage 2*, each run was composed of 39 S1 action potentials at a cycle length of 1000 ms and a final S2 stimulus delivered at an S1S2 interval determined by the APD_90_ of the final S1 beat, plus an offset of 10 ms, plus a diastolic interval (DI) with a range of 50-450 ms sampled using a Latin hyper-cube design with the other selected inputs as described below.

### Gaussian process emulators

Our overall approach is described in detail in a previous paper [18]. We treat the cardiac cell models as simulators which produce a vector of model outputs **y**_**s**_ as a function of a vector of model inputs (parameters) **x** such that **y**_**s**_ = *f*_*s*_(**x**). An emulator is then a statistical model of the simulator, sometimes known as a meta-model, a surrogate model, or a response surface model. The emulator approximates the model as **y**_**e**_ = *f*_*e*_(**x**), where the emulator output approximates the simulator output **y**_**e**_ ≈ **y**_**s**_ for a given input **x**.

In the present study we specified the emulator as a Gaussian process (GP), where the GP hyperparameters are optimized using a set of simulator runs called *design data*. When the GP has been fitted, the posterior prediction **y**_**e**_ at an input **x**^∗^ can be evaluated, which is a probability density with an expectation and a variance. The variance for the prediction **y**_**e**_ expresses uncertainty in the prediction of the simulator behaviour at **x**^∗^ [27, 28].

### Simulator runs for emulator design data

For *Stage 1* we generated design data from 300 runs of each cell model implemented as described above. The inputs for this design were sampled using an optimized Latin hyper-cube design ranging from ±50% of the central values (model defaults) given in Table 1 (i.e. from central value ×0.5 to central value ×1.5), except for *G*_*K*1_, *C*_*m*_, and extracellular ion concentrations. To reduce the incidence of instability (see below), these inputs had ranges of ±25%, ±25%, and ±10% respectively. A set of output biomarkers was obtained from the final action potential. A further set of 150 model model runs were then used for emulator validation (see below). For *Stage 2*, a second set of design data was produced from 200 simulator runs of each model, with a reduced set of inputs sampled from a Latin hyper-cube as described in the results section, and other inputs set to their central value. A further set of 100 simulator runs were used for emulator validation.

Outputs from the model runs used for *Stage 1* design data in the Courtemanche model and Maleckar model are shown in Fig 2. A wide variation in action potential shapes and durations were elicited by varying the model inputs.

**Fig 2.**
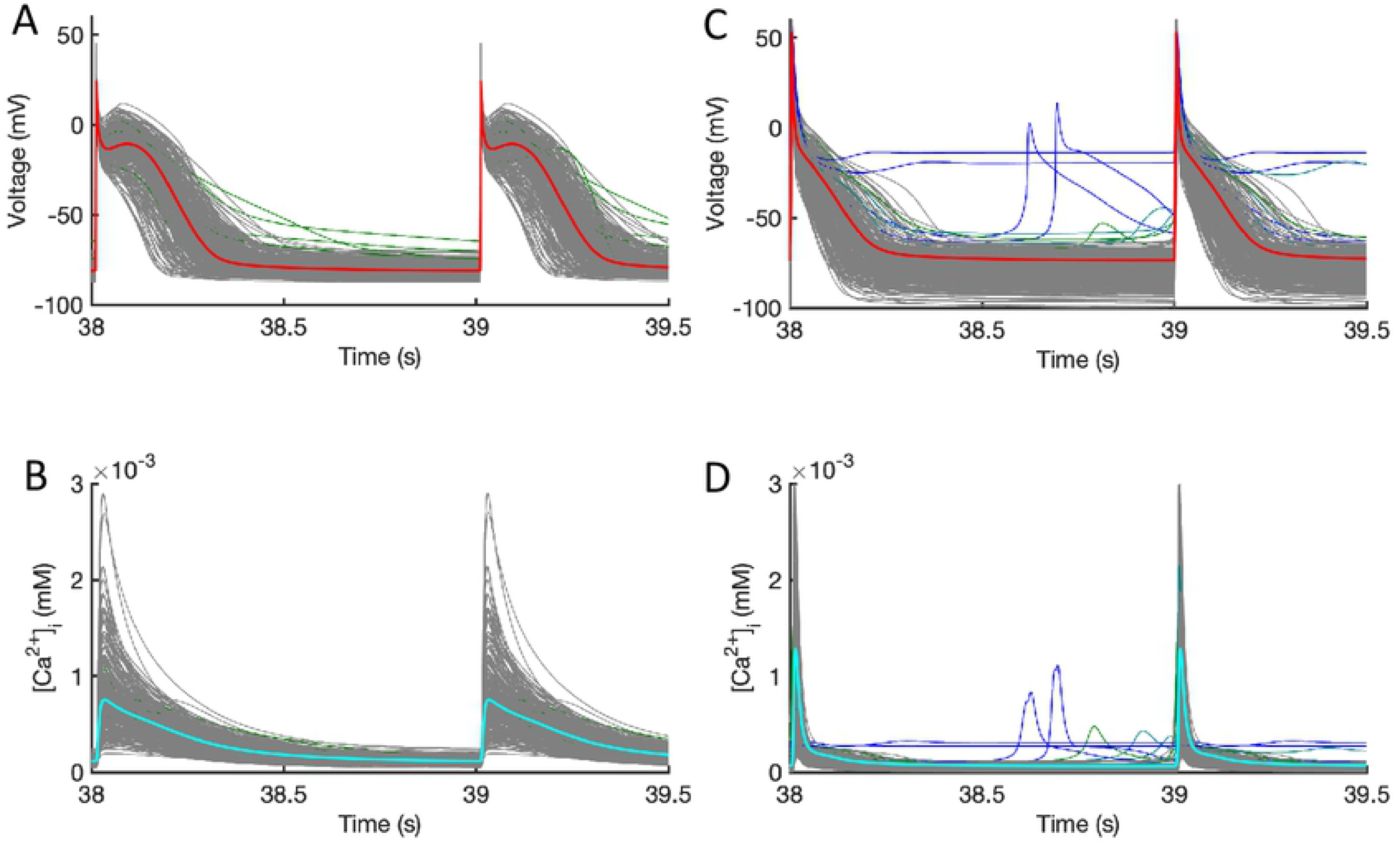
Design data. Action potentials and calcium transients produced by Latin hyper-cube sampling as described in the main text, and running each model with a pacing cycle length of 1000 ms. Traces shown in grey were used for emulator design data, and those shown in blue were excluded (see text for details). A, B: Courtemanche model. C, D: Maleckar model.

Model runs were removed from the design data if there was instability or abnormality in the model outputs evidenced by pacemaking activity, a resting potential greater than −60 mV, *APD*_90_ greater than 600 ms, or if *APD*_90_ of the 39th and 40th beats differed by more than 5% indicating alternans. Using these criteria, 5 out of 300 Courtemanche model runs in *Stage 1* were removed, because of a long APD or APD alternans. In the Maleckar model 14 out of 300 model runs in *Stage 1* were removed, all with pacemaking activity or failure to repolarise. In Fig 2 the removed runs are highlighted in blue. In *Stage 2* no model runs were removed for the Courtemenache model, and 5 model runs were removed for the Maleckar model.

### Emulator fitting

Our approach to fitting and using GP emulators is described in full detail elsewhere [18, 19]. Mathematical details including the expression for the posterior prediction of the emulator are provided in Supporting Information as well as in [27–29], and the Python code used in this study for emulator fitting, validation, sensitivity and uncertainty analysis is available from https://github.com/samcoveney/maGPy. Briefly, each emulator was composed of a mean function and a zero mean Gaussian Process,

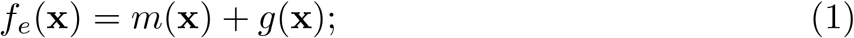

with a linear mean,

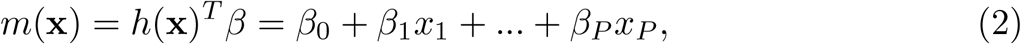

and a zero-mean GP,

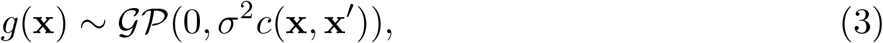

where the covariance has a Radial Basis Function form,

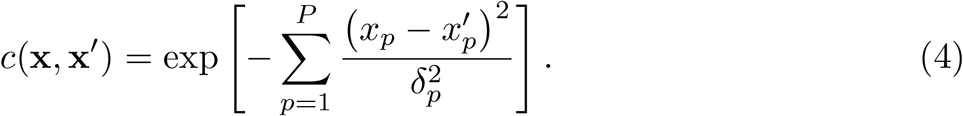

In these expressions **x**= (*x*_1_, *x*_2_, …, *x*_*P*_) are the *P* inputs (parameters), the emulator hyperparameters *β* and *δ* are vectors of length P, and *σ*^2^ is a scalar. The hyperparameters were obtained by maximum log-likelihood fitting to model inputs and outputs in the design data, assuming weak prior information on *β* and *σ*^2^ [29], and with a fixed nugget of 10^−7^ [30]. To avoid the fitting process becoming trapped in a local maximum, we repeated each fit ten times, each with a different set of randomly chosen initial values for the hyperparameters. The fit with the greatest log-likelihood was then selected. We produced a separate emulator for each of the outputs shown in Fig 1.

### Sensitivity and uncertainty analysis

Sensitivity and uncertainty analysis can be seen as distinct but related topics; where sensitivity analysis identifies the contribution of variance in each input to variance in each output, and uncertainty analysis concentrates on estimating the uncertainty in model outputs [31].

We calculated a first order sensitivity index for each combination of input and output [28]. For each input *i*, the first order sensitivity index describes how much the output variance would be reduced if *x*_*i*_ is fixed, while all other inputs are uncertain. The first order index is expressed as a proportion of the output variance calculated when all inputs are considered uncertain.

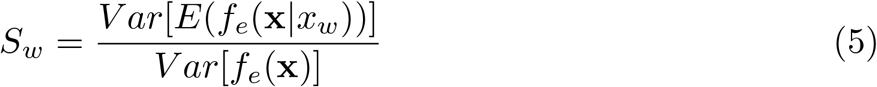

To capture the effect of interactions between the inputs, the total effect index can be calculated. This describes the reduction in output variance when *x*_*w*_ is uncertain and all other inputs are fixed, denoted as *x*_*~w*_. It is also expressed as a proportion of the output variance when all inputs are considered uncertain.

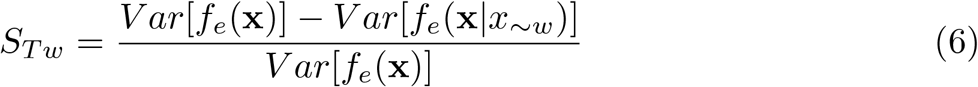

The difference between *S*_*Tw*_ and *S*_*w*_ is then the contribution of all interactions between *x*_*w*_ and *x*_*~w*_ to the variance in the output. These quantities were calculated using expressions given in the Supporting Information and described in [28]. Each uncertain input was assigned a mean of 0.5 in normalised units defined by the input ranges given in Table 1. Inputs that varied *±*50% were assigned a variance of 0.02, *G*_*K*1_ and *C*_*m*_ were assigned a variance of 0.04, and the extracellular ionic concentrations were assigned a variance of 0.1. We also calculated the main effect of each input on each output, using the procedure described in the Supporting Information, and the gradient of the main effect of a particular input around 0.5 normalised units was used to allocate a sign to the corresponding first order sensitivity index.

Several recent studies have calculated sensitivity indices based on partial least squares (PLS) regression [32, 33]. In this approach, each model output is assumed to be a weighted sum of inputs. Thus the model is described by the linear relationship

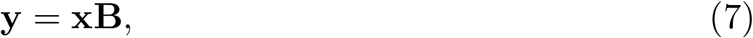

where **y** = (*y*_1_, *y*_2_, …, *y*_*M*_) is a vector of *M* outputs, **x** = (*x*_1_, *x*_2_, …, *x*_*P*_) a vector of *P* inputs, and **B** a *P* × *M* matrix of regression coefficients. An estimate of the matrix **B**, **B**_**PLS**_, can be found by PLS regression on a set of design data obtained from *N* model runs, that generates an *N* × *M* matrix of inputs **Y** and an *N* × *P* matrix of outputs **X**. Each element of **X**, *x*_*i,j*_ is regularised by subtracting the mean 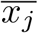 and dividing by the standard deviation of *x*_*j*_, and each element of **Y** is regularised in the same way. **B**_**PLS**_ is found by minimising the difference ‖**Ŷ** − **Y** ‖, where **Ŷ** = **XB**_**PLS**_.

The matrix **B**_**PLS**_ can be interpreted as a matrix of sensitivity indices, provided the linear model holds. Each element of **B**_**PLS**_, *b*_*i,j*_ describes how changing input *x*_*i*_ results in a corresponding change in output *y*_*j*_. In both cases the change is relative to the mean value, and is a fraction of its standard deviation.

For comparison with variance based sensitivity indices, we calculated PLS sensitivity indices from the *Stage 1* design data used to train the GP emulators. The input and output matrices **X** and **Y** were constructed from regularised design data inputs and outputs. The regression matrix **B**_**PLS**_ was then calculated using the Matlab function mvregress.

### Emulator validation

Each emulator was validated against an independent set of 150 simulator runs for *Stage 1* and 100 simulator runs for *Stage 2*. For each output, we calculated the mean average predicted error (MAPE) and the median individual standard error (ISE) for each validation run. The MAPE was given by

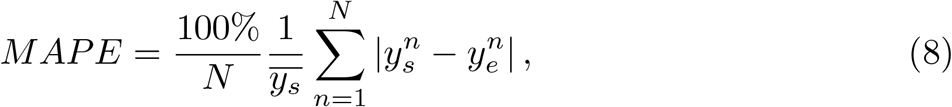

where N was the number of validation runs, 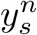 simulator output for run *n*, and 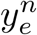 the posterior mean emulator output for run *n*. We used the mean of the simulator output 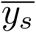 as a denominator instead of 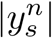 to avoid bias associated with small values of 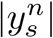.

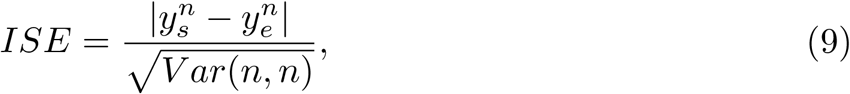

where 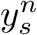 was the simulator output for run *n*, 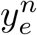 the posterior mean emulator output for run *n*, and *V ar*(*n, n*) the posterior emulator variance for run *n*.

For most stage *Stage 1* and *Stage 2* emulators the MAPE was less than 10% and the median ISE was less than 1.0. Most of the differences between the output from the emulator and the output of the simulator for a given set of inputs were small, and so the fit of the emulators was considered acceptable. A Table of MAPE and ISE is provided in the Supporting Information.

## Results

### *Stage 1* sensitivity indices

The first order sensitivity indices for both cell models are shown in Fig 3. Each row of the figure corresponds to one of the model outputs, and each column represents a model output. The sign of each sensitivity index was determined from the slope of the main effect (see below). The sum of the absolute values of the sensitivity indices for each output is given to the right of each grid. Since first order sensitivity indices are a proportion, a sum close to one indicates that almost all of the output variance is accounted for by the variance on each input. Smaller values for this sum, such as those for *APD*_50_ and *APD*_90_, can indicate that there are interactions among the inputs.

**Fig 3.**
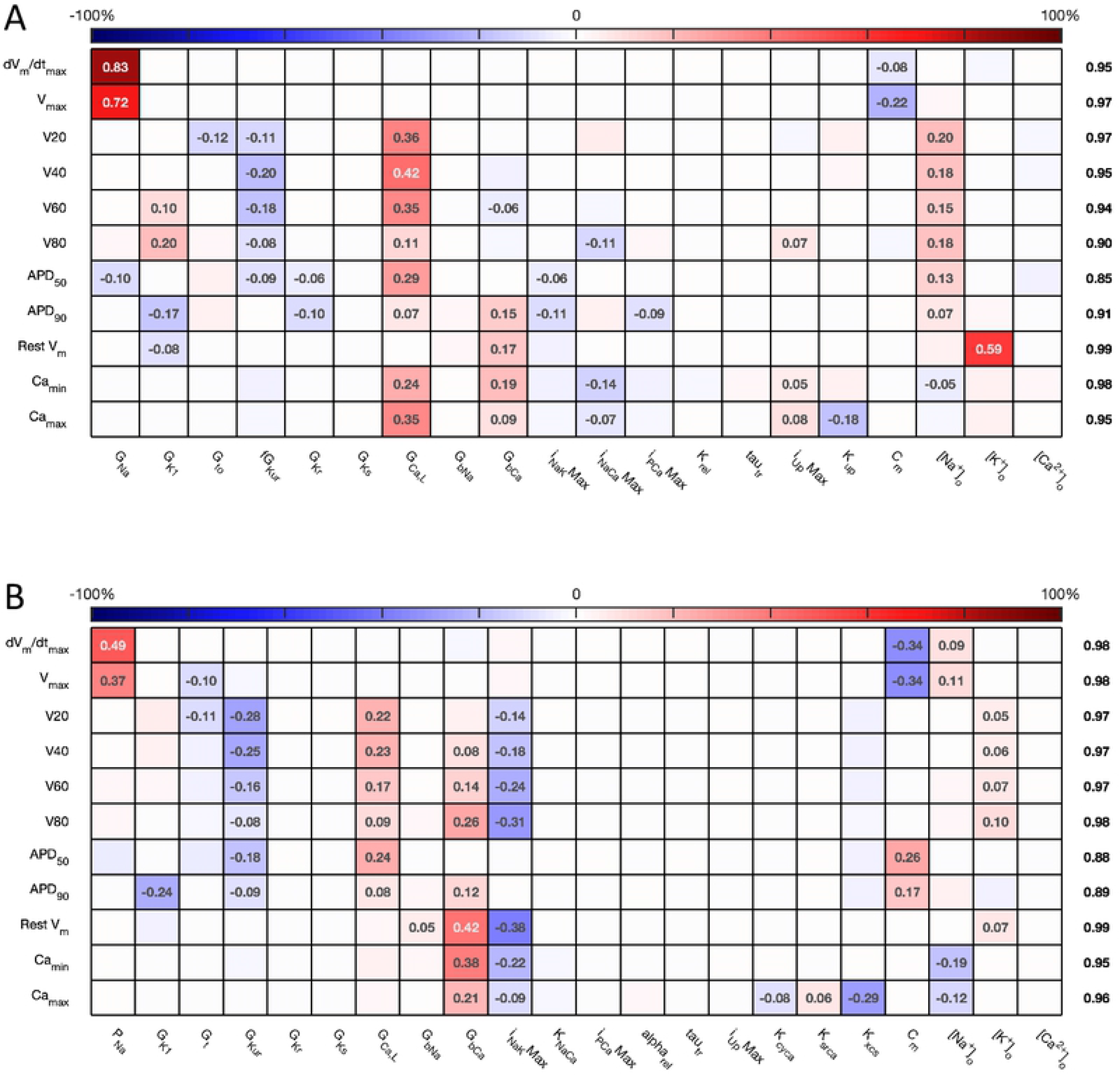
*Stage 1* first order sensitivity indices. The first order sensitivity index for each input and output is given, with a sign based on the gradient of the mean effect. Sensitivity indices less than 0.05 not shown, to assist visualization. The numbers at the right hand side of the table indicate the sum of the absolute values of sensitivity indices along each row. A: Courtemanche model. B: Maleckar model.

The total effect indices are shown in Fig 4. For each combination of input and output, the difference between the total effect index and the first order index reflects interactions with the other inputs. The sum of these differences is shown at the right hand side of Fig 4. In most cases the first order and total effect indices are similar, and the sum of differences is small indicating that the effect of interactions is small. However, the sum of differences is larger for *V*_80_ in the Courtemanche model, as well as for *APD*_50_ and *APD*_90_ in both models, and we conclude that there are interactions between inputs, and these interactions have an effect on repolarisation.

**Fig 4.**
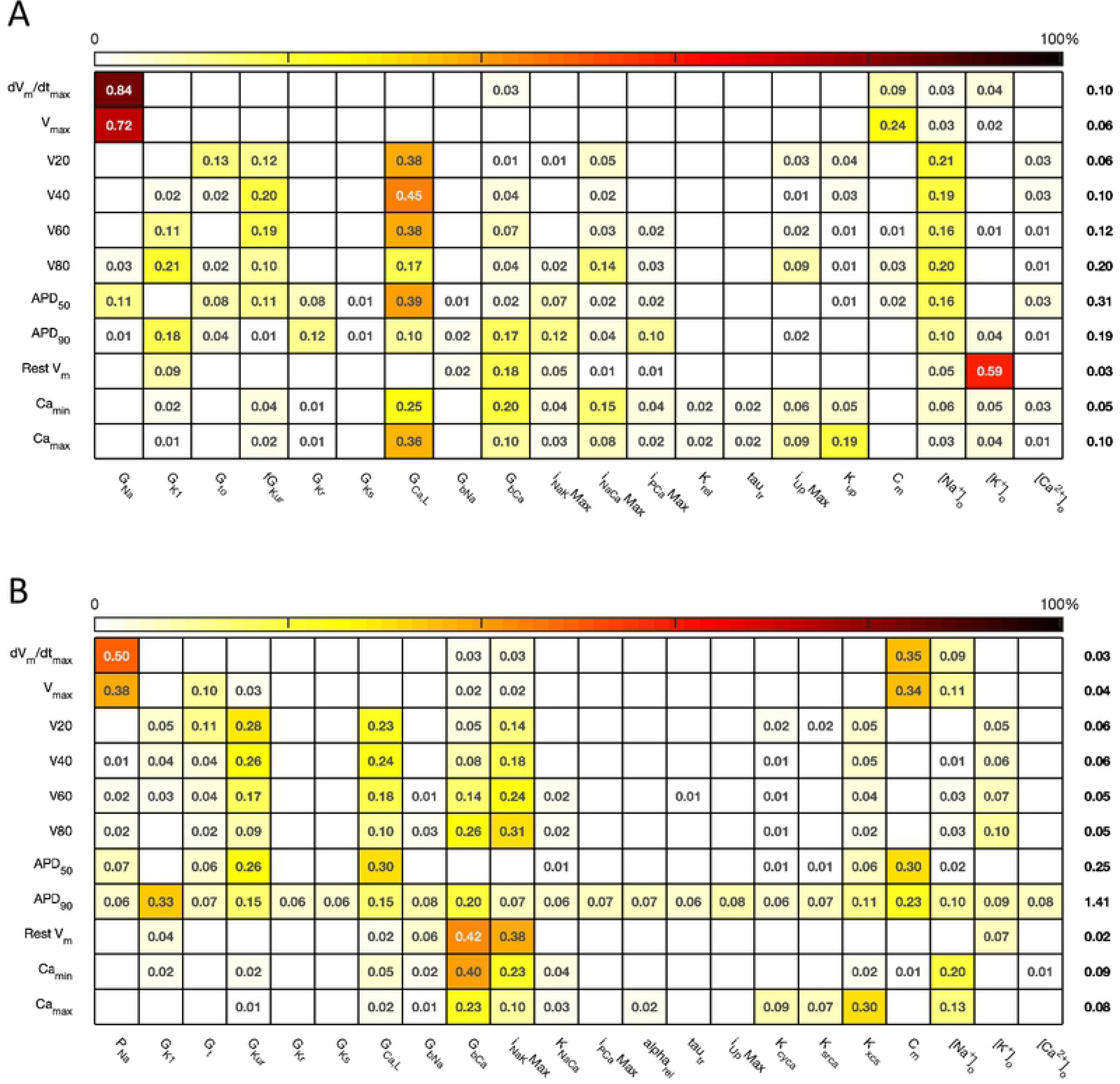
*Stage 1* total effect indices. The total effect sensitivity index for each input and output is given, indices less than 0.01 not shown. Numbers on right hand side are the the sum across each row of the differences between the total effect index and the absolute value of the first order index. A: Courtemanche model. B: Maleckar model.

For comparison, Fig 5 shows sensitivity indices obtained by PLS regression on the emulator design data, and a comparison with variance based first order indices. The comparison plots show broad agreement, with the first order indices 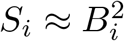. This relationship arises from the different definitions of *S*_*i*_ and *B*_*i*_ based on variance and standard deviation respectively [34].

**Fig 5.**
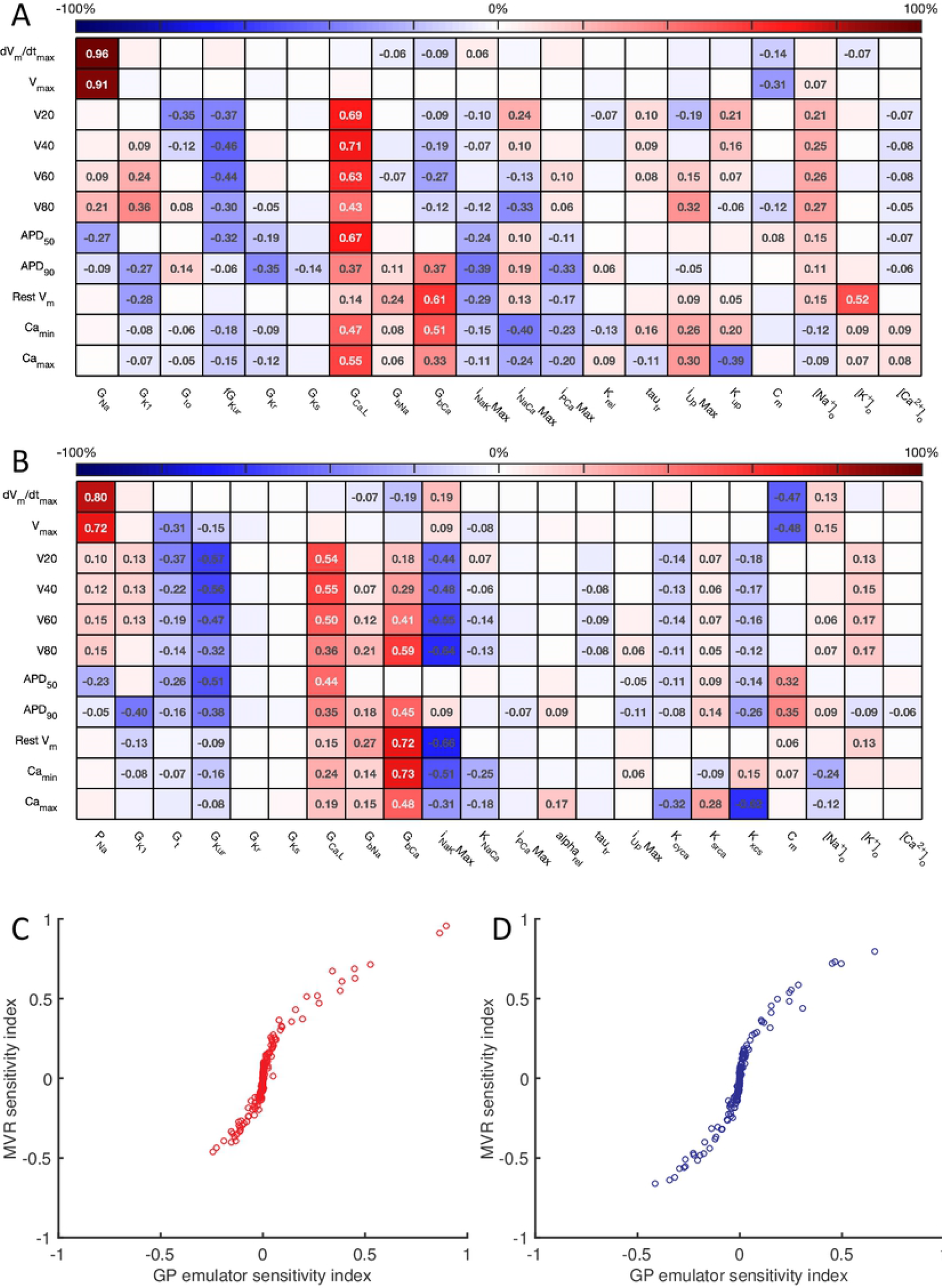
Multivariate regression sensitivity indices. The sensitivity index based on multivariate regression for each input and output is given. Right hand panels compare these sensitivity indices with those obtained from GP regression and shown in Fig 3. A: Courtemanche model. B: Maleckar model.

Overall these sensitivity indices show the contribution of uncertainty in each input to uncertainty in each output. Thus the main contributors to uncertainty in *dV/dt*_*max*_ are the *Na*^+^ channel maximum conductances *G*_*Na*_ and *P*_*Na*_, and the membrane capacitance *C*_*m*_. The sign of the sensitivity indices show that these act in opposite directions, as would be expected from the role played by the *Na*^+^ current in depolarisation: increasing *Na*^+^ current acts to increase *dV/dt*_*max*_, whereas increasing *C*_*m*_ acts to decrease *dV/dt*_*max*_. The bigger influence of *C*_*m*_ in the Maleckar model arises becuase the stimulus current scales with 1/*C*_*m*_; a larger *C*_*m*_ results in a smaller stimulus current, which in turn produces a smaller *dV/dt*_*max*_ and a smaller *V*_*max*_.

In both models, these sensitivity indices can be interpreted to show that increased outward currents (such as *G*_*Kur*_) act to decrease both the voltage of the action potential plateau and action potential duration, whereas increased outward currents *G*_*GaL*_ and *G*_*bCa*_ have an opposite effect. In the Maleckar model *I_NaK_Max* has the opposite effect to *G*_*bCa*_. In the Courtemanche model, 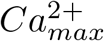 and 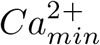 are influenced by *G*_*CaL*_, *G*_*bCa*_, *I*_*NaCaMax*_ and *Ca*^2+^ handling parameters *I*_*rel*_ and *K*_*up*_, whereas in the Maleckar model *G*_*CaL*_ and *K*_*NaCa*_ have a negligible effect, and this reflects the different formulation of *Ca*^2+^ handling in the two models [7].

The sensitivity analysis shows that several inputs influence *APD*_50_, *APD*_90_, *RestV*_*m*_, and 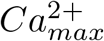. Fig 6 shows the main effect of each input on these outputs, for each cell model. The main effect shows the expected value of the output as each input is fixed and varied in turn across the normalised range 0 … 1 corresponding to the input ranges given in Table 1, while all other inputs are considered uncertain. The residual variance arising from the uncertain inputs accounts for the fact that the main effects do not converge exactly for an input value of 0.5.

**Fig 6.**
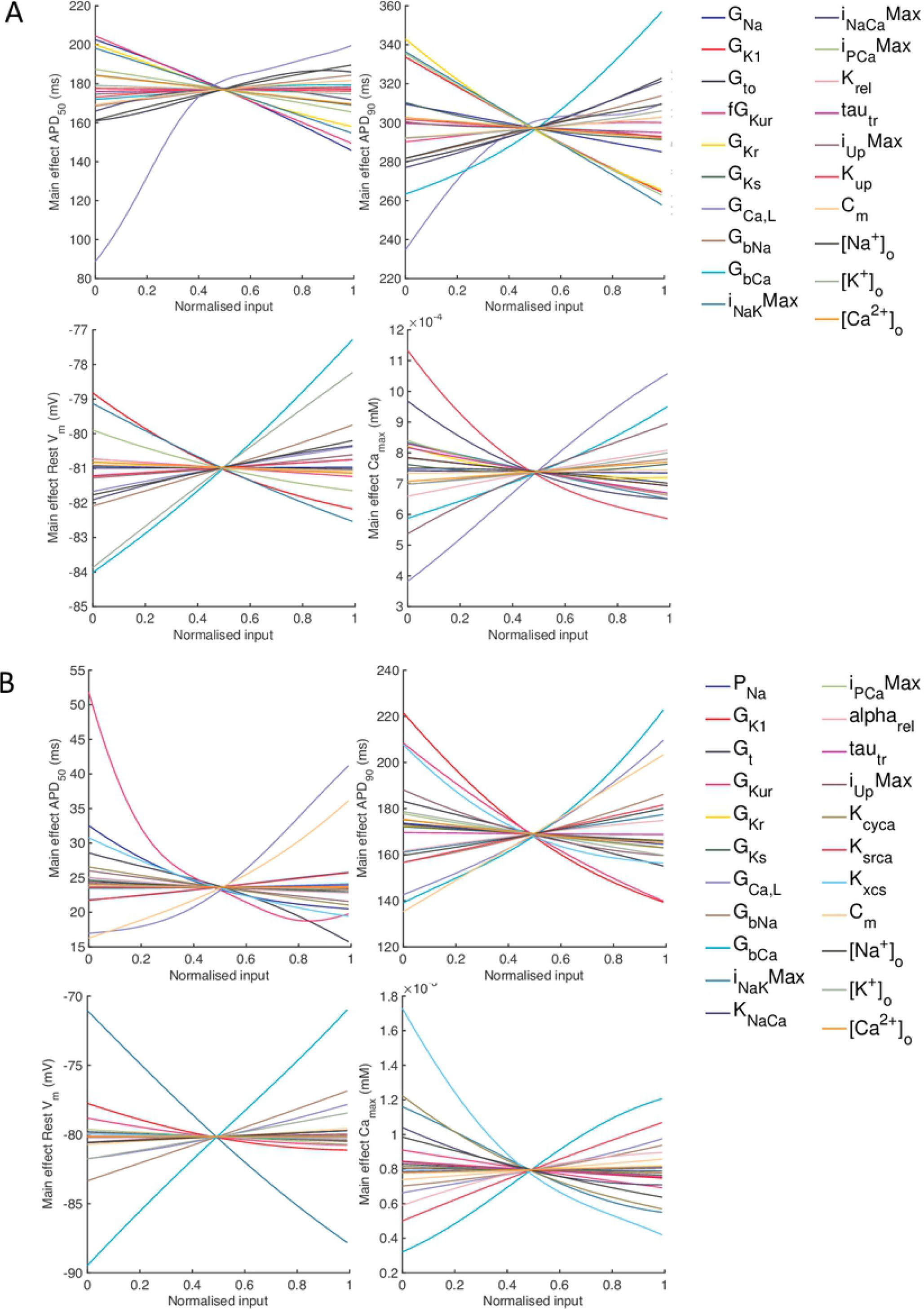
Stage 1 selected main effects plots. Main effects for *APD*_50_, *APD*_90_, *RestV*_*m*_, and 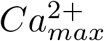. A: Courtemanche model. B: Maleckar model.

Some of the effects are comparable between the two models, for example increasing *G*_*bCa*_ acts to increase *APD*_90_, *RestV*_*m*_, and 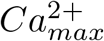. Several of the effects are nonlinear, for example the main effect of *G*_*CaL*_ on *APD*_50_ and *APD*_90_. However, the overall picture is complex, and it is hard to compare the different models. In order to simplify the analysis, we selected a subset of inputs for *Stage 2* of the analysis based on their sensitivity indices as described below.

### *Stage 2* sensitivity analysis

For *Stage 2*, we concentrated on inputs that strongly influenced action potential shape and duration, with first order sensitivity index of more than 0.1. To simplify the analysis further, we excluded *G*_*Na*_ and *P*_*Na*_ as these inputs mainly influence action potential upstroke and amplitude. We also excluded extracellular concentrations, since these are tightly controlled in normal physiological conditions, and we excluded the inputs directly involved in the storage, uptake and release of intracellular *Ca*^2+^ because we sought to concentrate on action potential shape and duration. The *Stage 2* inputs selected for the Courtemanche model were *G*_*K*1_, *G*_*to*_, *G*_*KurMult*_, *G*_*CaL*_, *G*_*bCa*_, *I*_*NaK*_*Max*, *I*_*NaCa*_*Max*, and *I*_*PCa*_*Max*, and for the Maleckar model *G*_*K*1_, *G*_*t*_, *G*_*Kur*_, *G*_*CaL*_, *G*_*bCa*_, *I*_*NaK*_*Max*, and *C*_*m*_. All other inputs were assigned their central value from Table 1. In addition, the DI of the final action potential was introduced as an additional input to explore the dynamic behaviour of the model.

The sensitivity indices for *Stage 2* are shown in Fig 7 and 8, and the main effects for *APD*_50_, *APD*_90_, *RestV*_*m*_, and 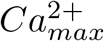 in Fig 9.

**Fig 7.**
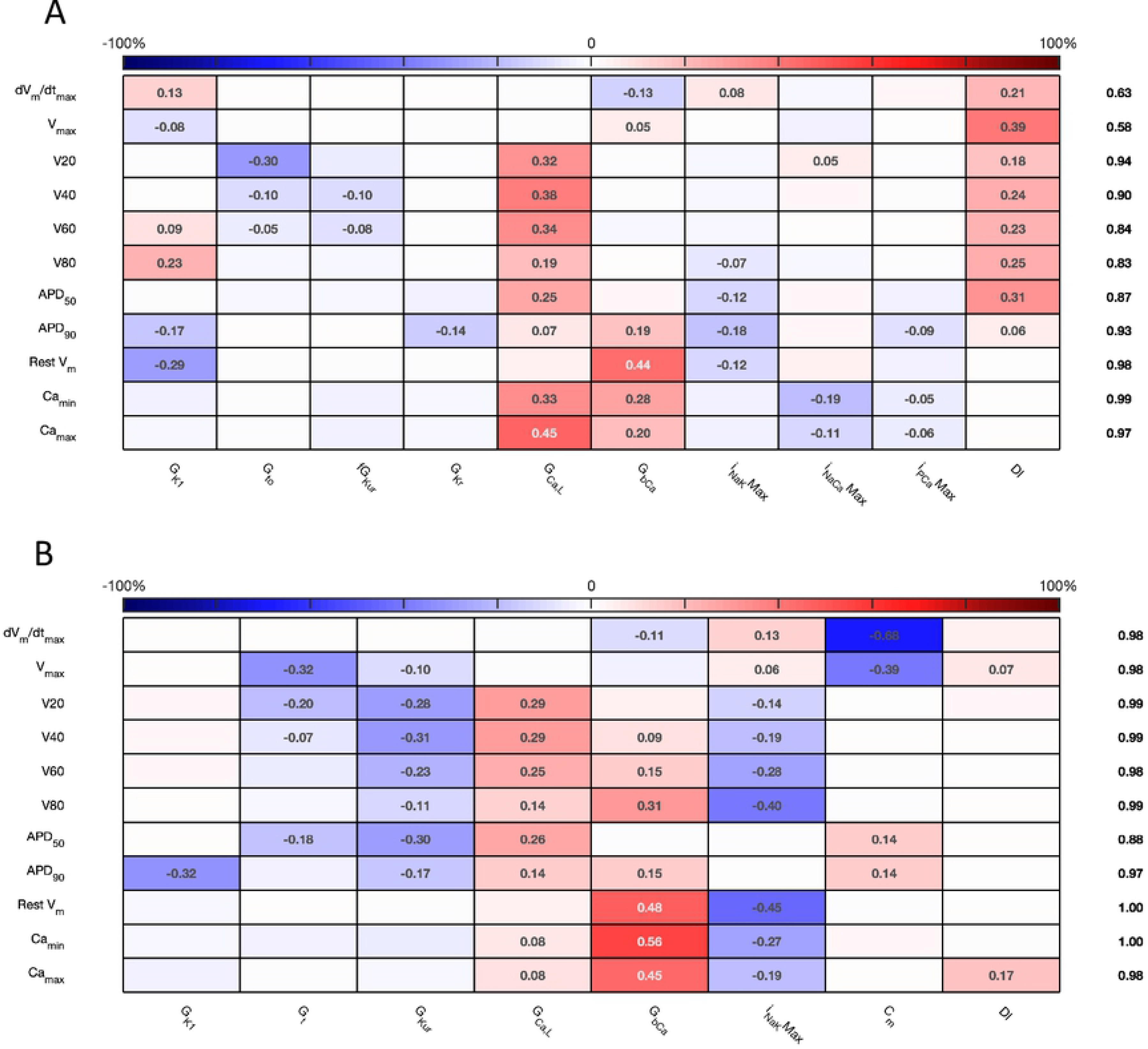
*Stage 2* first order sensitivity indices. The first order sensitivity index for each input and output is given, with a sign based on the gradient of the mean effect. Sensitivity indices less than 0.05 not shown. The numbers at the right hand side of the table indicate the sum of sensitivity indices along each row. A: Courtemanche model. B: Maleckar model.

**Fig 8.**
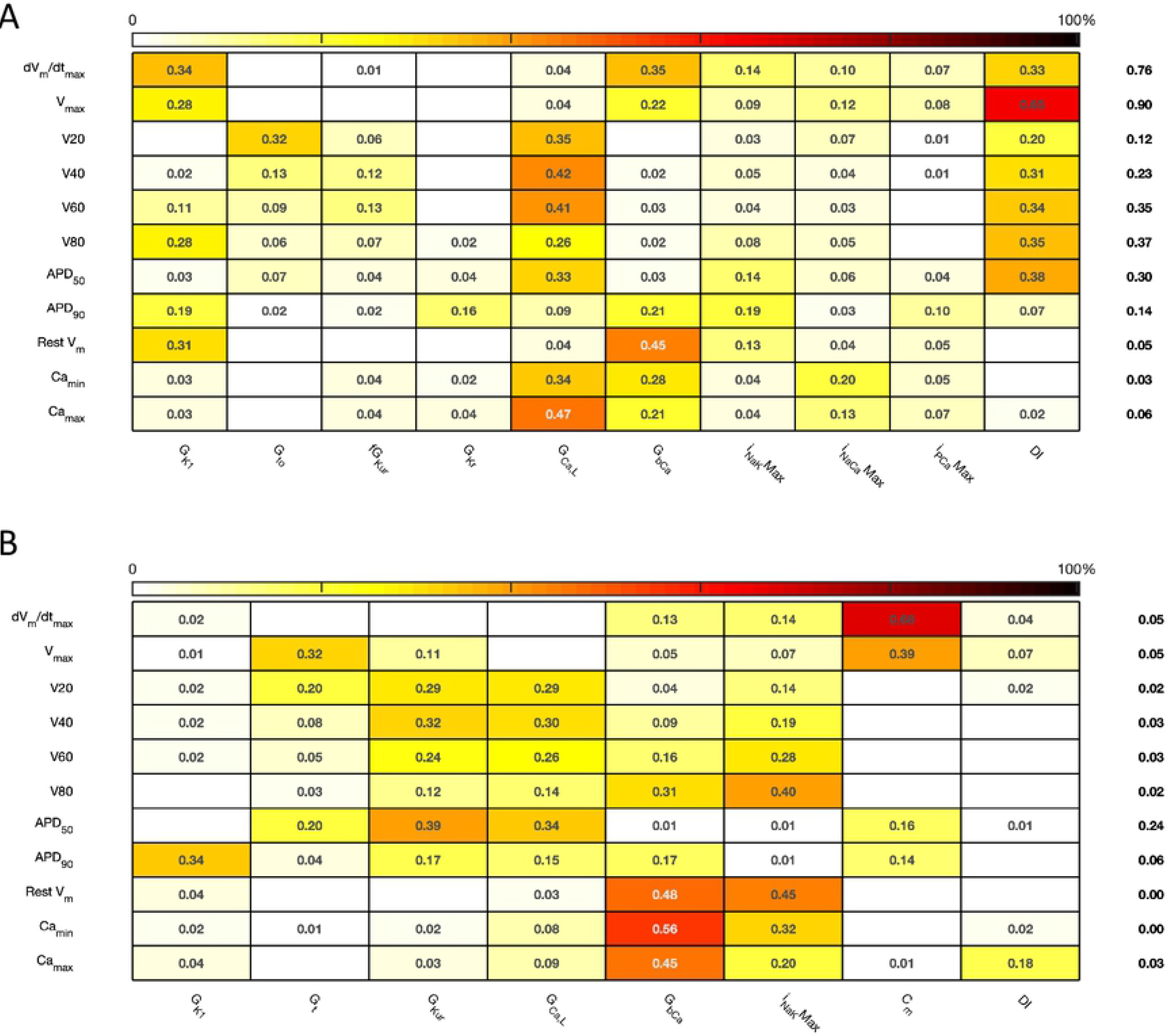
*Stage 2* total effect sensitivity indices. The total effect index for each input and output is given, indices less than 0.01 not shown. A: Courtemanche model. B: Maleckar model.

**Fig 9.**
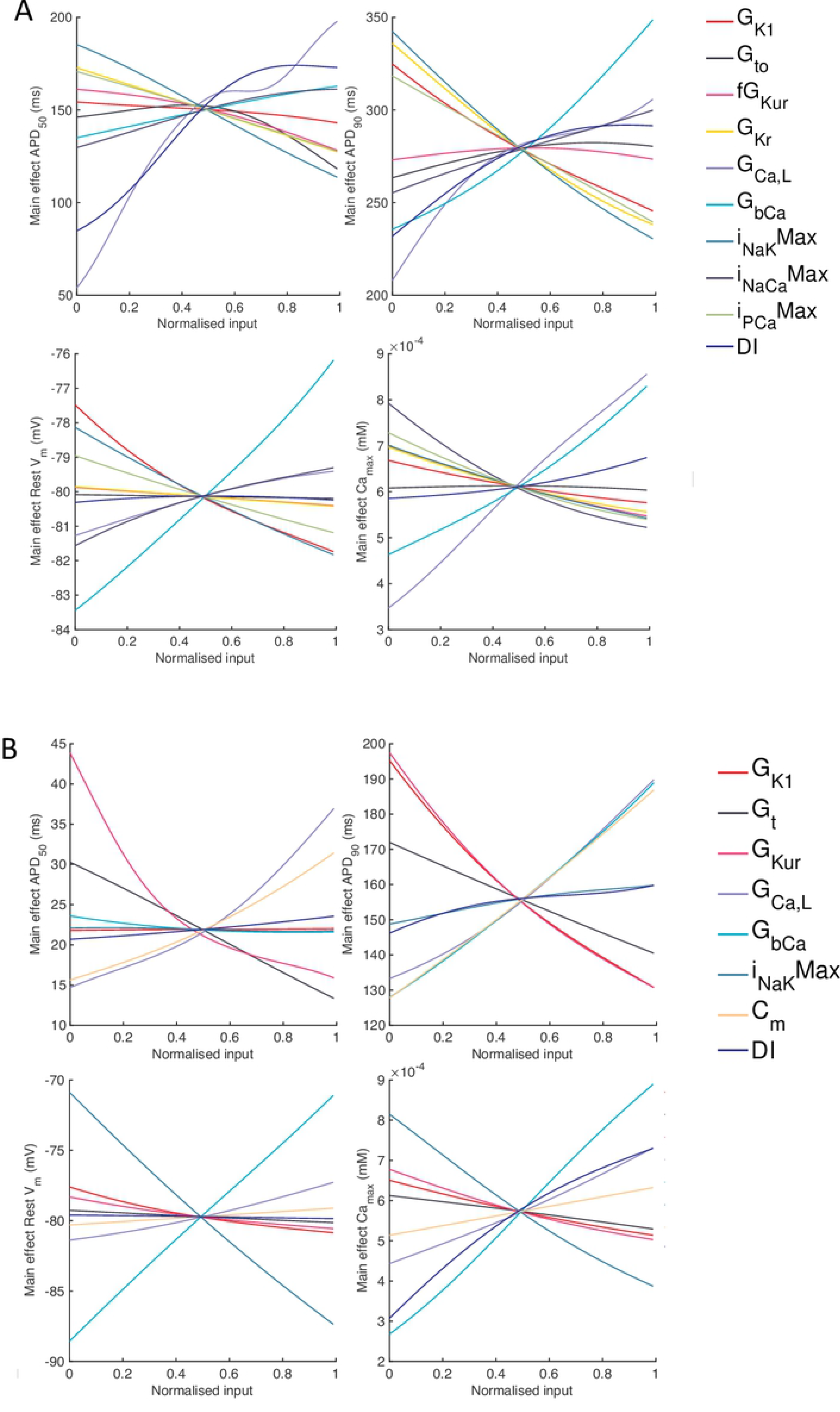
*Stage 2* selected main effects plots. Stage 2 main effects for *APD*_50_, *APD*_90_, *RestV*_*m*_, and 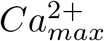. A: Courtemanche model. B: Maleckar model.

Overall, both first order and total effect sensitivity indices were similar to *Stage 1*, but the interactions for the Courtemanche model were larger than for *Stage 1*. In the Courtemanche model both first order and total effect indices for *DI* were larger than for the Maeckar model, indicating that *DI* acts to influence both the shape and duration of the action potential in the Courtemanche model.

The main effects plots show opposite effects of inward and outward currents on action potential duration; however *APD*_50_ as a proportion of *APD*_90_ in the Maleckar model was considerably shorter than in the Courtemanche model as a result of the different action potential shape, and so both the sensitivity indices and main effects for *APD*_50_ may not be easily comparable. This observation is reflected in the larger main effect of *DI* on *APD*_50_ in the Courtemanche model. The main effects plots also show several nonlinear relationships, for example the main effect of *G*_*Kur*_ was nonlinear in each of the outputs shown in Fig 9. The main effect of *G*_*Kur*_ on *APD*_90_ in the Maleckar model was larger, and in an opposite direction, to the main effect in the Courtemanche model. This was explored in additional simulations where *G*_*Kur*_ was decreased to 50% and increased to 150% of its default value. Fig 10 shows a simulated action potential, together with the principal inward and outward currents that act during the action potential plateau.

**Fig 10.**
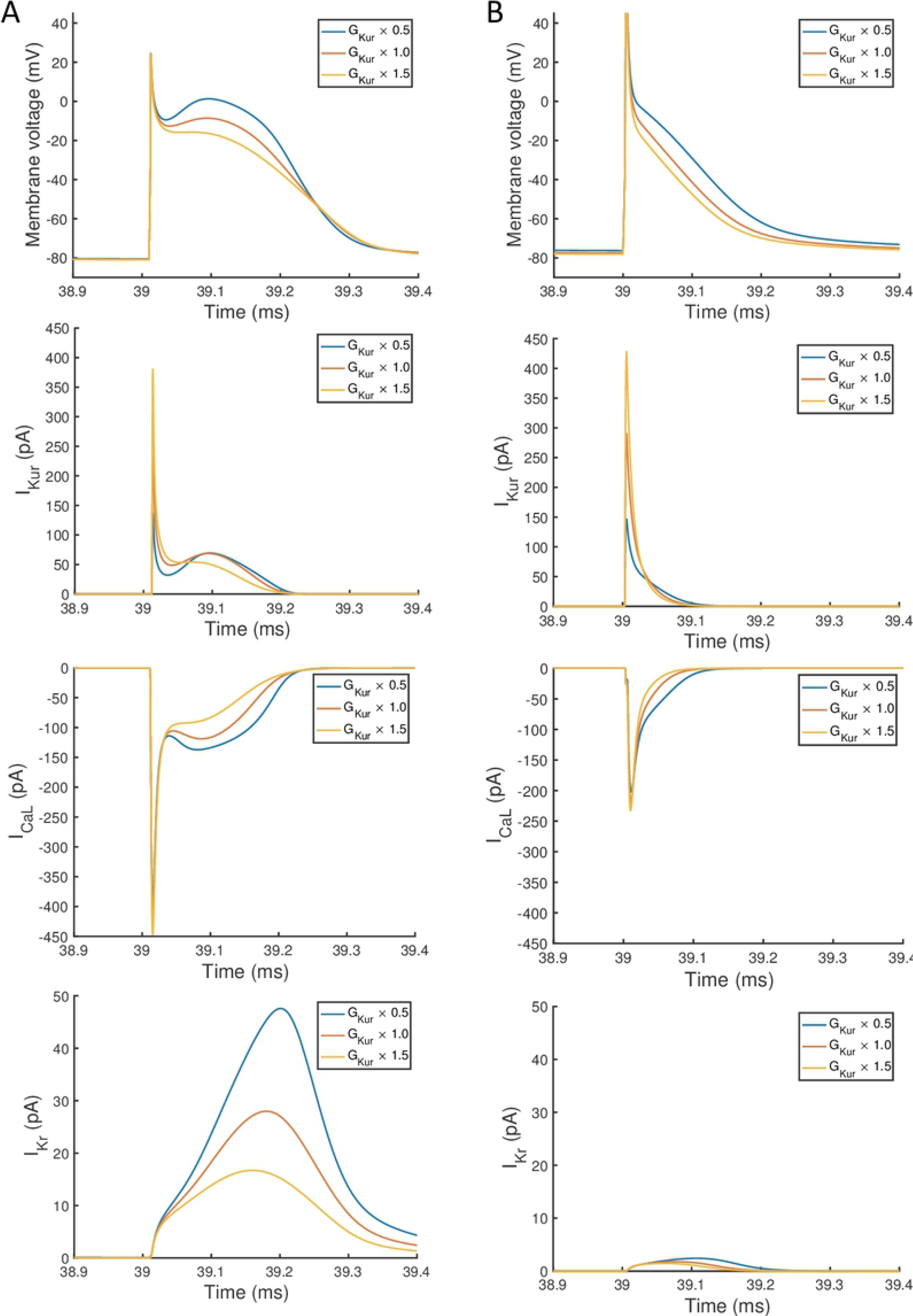
Effect of magnitude of *I*_*Kur*_ on inward and outward currents. Simulated action potential, outward current *I*_*Kur*_, inward current *I*_*Ca,L*_, and outward current *I*_*Kr*_. In each case the final beat of 40 is shown, with a pacing cycle length of 1000 ms. A: Courtemanche model. B: Maleckar model.

Changing *G*_*Kur*_ influenced the voltage of the action potential plateau in both models, but had a different effect on the timing of repolarisation because of different current formulations in each model. The time course of *I*_*to*_ in both models was similar, and was not strongly influenced by *G*_*Kur*_ and so is not shown. A decrease in *G*_*Kur*_ reduced the outward current during the initial part of the action potential plateau. This resulted in an increased voltage during the plateau, and a larger inward *I*_*Ca,L*_, which is voltage-dependent and acts to prolong the action potential plateau. In turn, the increased plateau voltage resulted in greater activation of the outward current *I*_*Kr*_, which is larger in the Courtemanche model compared to the Maleckar model. Thus in the Courtemanche model, a decrease in *G*_*Kur*_ resulted in increased *I*_*Kr*_, with little change in action potential duration. In the Maleckar model *I*_*Kr*_ is much smaller, and so the increased plateau voltage did not result in increased outward current during repolarisation, and so action potential duration was prolonged.

### APD restitution

The *Stage 2* analysis included diastolic interval as an input, which enabled us use the emulators to examine how different inputs affect APD restitution. In Fig 11 we have plotted a surface showing the expected value of *APD*_90_, coloured by the 95% credible interval (see Supporting Information). In each of these plots the emulators were evaluated with all inputs assigned fixed values with no uncertainty, and so the 95% credible intervals reflect only uncertainty in the emulator fit, with no uncertainty arising from uncertainty in the inputs. We assigned a value of 0.5 in normalised units to all of the inputs, except for *DI* and another inputs that were varied in each plot; these were assigned fixed values between 0 … 1.

**Fig 11.**
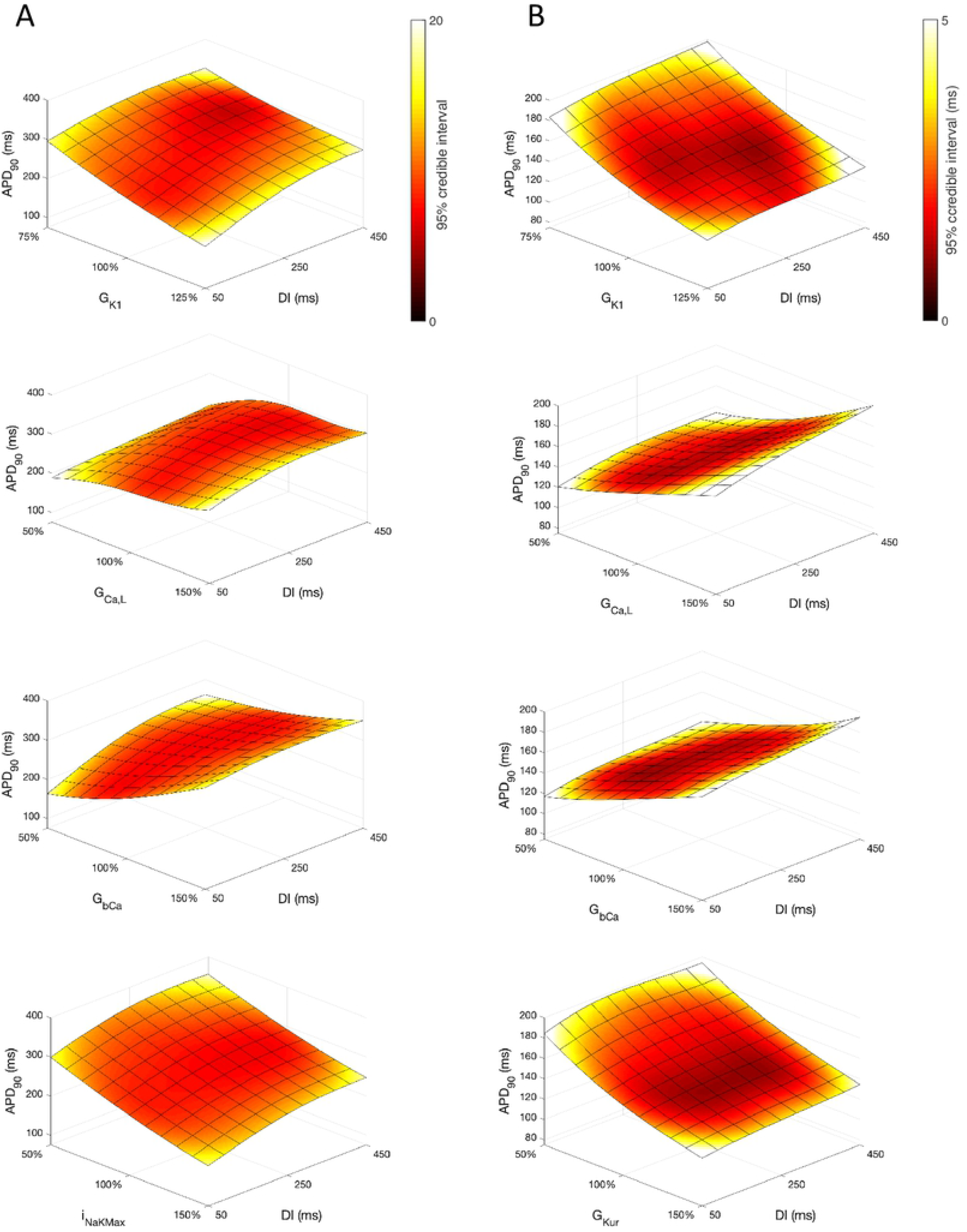
*APD*_90_ as a function of *DI* and other inputs. Each plot shows a surface of predicted mean *APD*_90_, coloured by the 95% credible interval. A: Courtemanche model. B: Maleckar model.

Overall, *APD*_90_ restitution was flatter in the Maleckar model compared to the Courtemanche model, which is consistent with other studies [6, 7]. The overall effect of *G*_*K*1_, *G*_*CaL*_ and *G*_*bCa*_ on *APD*_90_ was similar in each model, with *I*_*NaKax*_ in the Courtemanche model having a similar effect to *G*_*Kur*_ in the Maleckar model.

However the shape of the *APD*_90_ restitution was modulated to some extent by the other inputs shown. In the Courtemanche model, decreasing *G*_*bCa*_ resulted in steepening of *APD*_90_ restitution, with a marked decrease in *APD*_90_ for short *DI* as *G*_*bCa*_ was changed from 150% to 50% of its central value. The Maleckar model also showed a marked decrease in *APD*_90_ as *G*_*bCa*_ was reduced, but this change was seen across the full range of *DI*.

### Uncertainty analysis

We assessed how the uncertainty in model outputs changed as uncertainty in all of the model inputs was increased, and the results for *APD*_90_ are shown in Fig 12. Uncertainty in *APD*_90_ increased monotonically with uncertainty in the inputs, for both *Stage 1* and *Stage 2*. Reducing the number of uncertain inputs for *Stage 2* acted to reduce the output uncertainty for the Maleckar model, but increased output uncertainty slightly in the Courtemanche model. This rather unexpected finding may be due to interactions among the inputs. However, in the Maleckar model, the total effects for *APD*_90_ in Fig4B are higher than the total effects for the Courtemanche model, indicating a greater degree of interaction in the Maleckar than Courtemanche models. The smaller number of inputs for *Stage 2* could then result in less uncertainty arising from interactions among uncertain inputs for the Maleckar model, and a consequent reduction in uncertainty. However fixing some inputs for the *Stage 2* analysis may have had an effect on interactions the in the Courtemanche model, leading to an increase in uncertainty in predicted *APD*_90_. An alternative explanation would be that with some of the inputs fixed, the uncertain inputs contribute mote to output uncertainty in the Courtemanche model.

**Fig 12.**
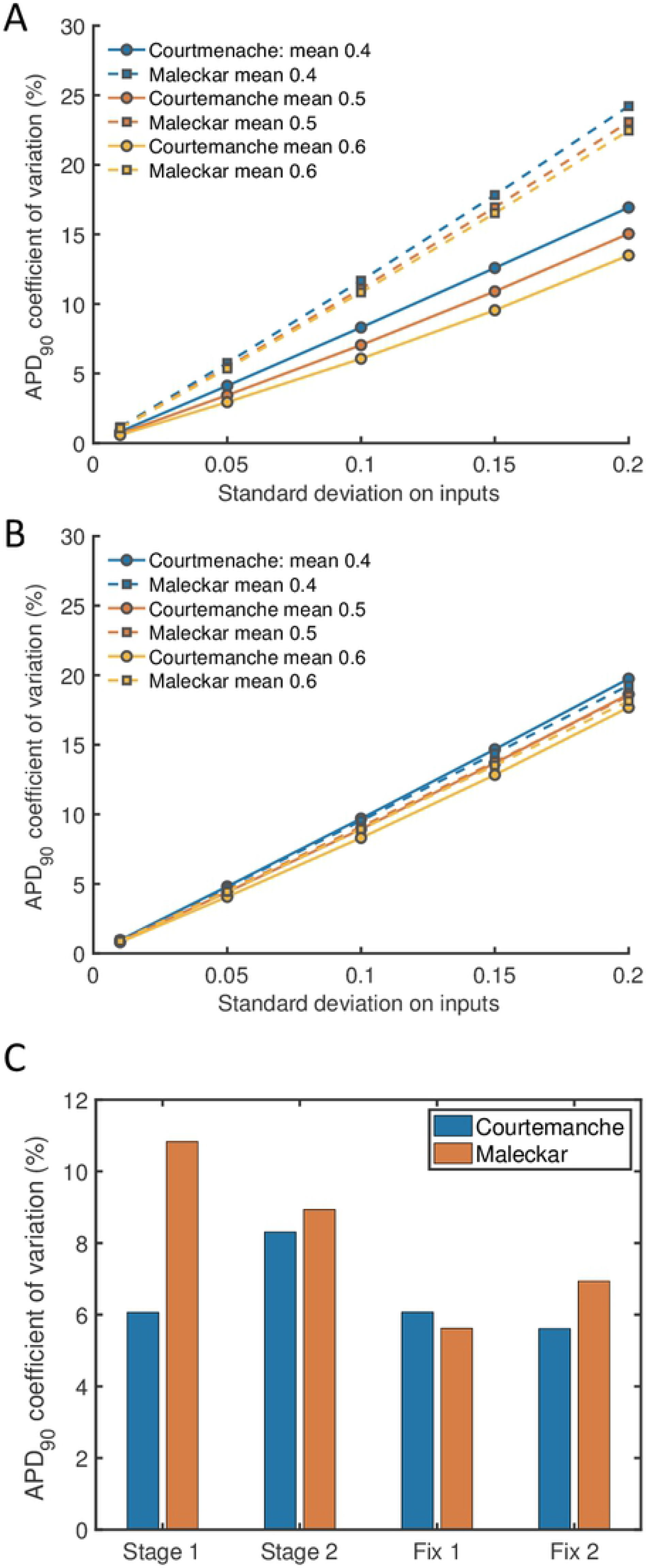
Uncertainty in *APD*_90_ and *V*_*max*_. Coefficient of variation (standard deviation divided by mean) in *APD*_90_ for *Stage 1* (A) and *Stage 2* plotted against imposed uncertainty on all inputs (B) for each model, with different mean values of 0.4, 0.5, and 0.6 in normalised units. (C) Coefficient of variation in *APD*_90_ for *Stage 1*; *Stage 2*; simulations where *G*_*K*1_, *G*_*Kr*_ (*G*_*Kur*_ for Maleckar model), and *G*_*bCa*_ were fixed at 0.5 and all other inputs were uncertain with standard deviation of 0.1 (Fix 1); and simulations where all other inputs were fixed with mean of 0.5, and *G*_*K*1_, *G*_*Kr*_ (*G*_*Kur*_ for Maleckar model), and *G*_*bCa*_ were uncertain with standard deviation of 0.1.

The first two bars of Fig 12C show the coefficient of variation for *APD*_90_ with inputs set to a mean of 0.5 and standard deviation of 0.1 in normalised units, corresponding to a single point for each model in Fig 12 A and B. The third set of bars (denoted Fix 1) show the coefficient of variation for *APD*_90_ when the three inputs with the highest sensitivity indices are fixed at 0.5, and the fourth set of bars (denoted Fix 2) show the coefficient of variation when all inputs except these three inputs are fixed at 0.5.

## Discussion

In this study we have obtained novel insights into the comparative mechanisms of two atrial cell models, and have demonstrated the use of Gaussian process emulators for sensitivity and uncertainty analysis.

### Human atrial cell models

Models of the human atrial action potential are a subject of research interest and clinical relevance because heterogeneity in action potential shape and duration in different parts of the atria is associated with vulnerability to atrial fibrillation [35], and persistence of atrial fibrillation is associated with remodelling of the atrial action potential [23, 36]. There are several different models of human atrial myocytes, each with different properties [6, 7, 37]. Most studies of these models have focussed on the mechanisms that change action potential duration because a reduced APD increases vulnerability to atrial fibrillation, and APD can be modulated pharmacologically [7, 9, 12, 21, 38]. These previous studies have highlighted the importance of *I*_*CaL*_, as well as *I*_*K*1_ and *I*_*Kur*_, in regulating APD. The present study adds to our understanding of the Courtemanche and Maleckar models by providing a more comprehensive view of how the model parameters affect the shape and duration of the action potential, as well as the maximum and minimum of the *Ca*^2+^ transient.

Increased inward current tends to increase amplitude of the upstroke and plateau as well as increasing APD, and increased outward current tends to have the opposite effect. The present study has highlighted how action potential duration and shape depends on the net flow of charge across the cell membrane, which is finely balanced so small changes in the magnitudes of inward and outward currents can strongly influence action potential shape and duration. The two models examined in this study represent repolarisation using a different balance of currents, and this difference along with the consequences can be seen in Fig 10. The relative magnitudes of *I*_*CaL*_ and *I*_*Kr*_ in the two models are different, with much smaller *I*_*CaL*_ and *I*_*Kr*_ in the Maleckar model. This may explain why *G*_*bCa*_ exerts a stronger influence over 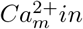 and 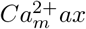 in the Maleckar model compared to the Courtemanche model.

Overall, the difference between the first order sensitivity indices (Fig 3) and the total effect indices (Fig 4) was small. The sum of these differences for each output (right hand column in Fig 4 and Fig 8) indicates more interactions in the Courtemanche model than in the Maleckar model, and that these interactions tend to affect the plateau of the action potential. These observations mean that overall the interactions between the inputs in these models do not have a strong effect on the outputs, and so we can conclude that the inputs examined in this study tend to act independently. This is a potentially important feature of the models, which could be exploited for model calibration as well as for examining mechanisms of remodelling and pharmacological action. However, it remains to be seen whether this independence is a feature of real cardiac myocytes.

### Sensitivity and uncertainty analysis

As models of cardiac cell and tissue electrophysiology become more widely used, it is becoming increasingly important to understand how different components of the models influence model behaviour, and especially how these different components interact. Biophysically detailed cardiac cell models are complex, with many interacting parts. Some of these model components may be inherited from earlier models and experiments [8], and the process by which model parameters are fitted is also fragile when there are uncertainties associated with experimental data [5]. The development and evaluation of tools for sensitivity and uncertainty analysis of cardiac models is therefore an important and growing area [16], but much remains to be done.

Several recent studies have pioneered the use sensitivity indices obtained by of partial least squares (PLS) regression of simulator outputs on simulator inputs, which allows a calculation of sensitivity indices [9, 10, 32]. This approach is straightforward to implement, and we have found that it gives sensitivity indices that agree well with the square root of the first order index obtained using the GP approach (Fig 5), and the reason for this appears to be that the PLS and GP indices are based on variance and standard deviation respectively. The overall agreement indicates that both approaches can yield similar first order sensitivity indices, although the GP easily enables calculation of interaction effects as well as first order indices. Other approaches for uncertainty and sensitivity analysis based on generalised polynomial chaos expansion have also been developed and used for analysis of cardiovascular system models [17]. These approaches also enable calculation of sensitivity indices, but the relative merits of these different approaches are only beginning to be explored [43]. Both GP emulators and polynomial chaos expansions enable the explicit treatment of uncertainties, and so offer advantages for more comprehensive model analysis.

### Gaussian process emulation of biophysical models

The present study has focussed on Gaussian process (GP) emulators, which are one class of tools for sensitivity and uncertainty analysis. A GP acts to interpolate an output surface, providing a probabilistic estimate of a simulator output for a particular point in a high dimensional input space. A GP is quick to evaluate. It can be trained on a relatively small number of simulator runs, and so offers advantages over other approaches that rely on large numbers of simulator evaluations [12, 39].

A GP is usually trained using a set of design data composed of a set of simulator inputs and outputs. The emulator hyperparameters are obtained conditional on the design data using a maximum likelihood approach. The quality of the emulator fit can then be assessed by comparing the outputs obtained from the emulator for a particular set of inputs with those obtained by the simulator for the same set of inputs. In the present study we used the MAPE and ISE to quantify the difference between emulator and simulator outputs, although other measures such as the Mahanalobis distance can be used [18]. These measures provide some guidance about whether the emulator is under or over-fitted to the design data.

The number of simulator runs required to compose the design data remains an open question, and will depend on the complexity of the model output surfaces. A typical rule of thumb is to use ten times the number of inputs, and based on our previous experience [18] we opted for 300 runs for *Stage 1* and 200 runs for *Stage 2*. We also trained the *Stage 1* emulators on design data sets composed of 200 and 400 simulator runs, and obtained similar sensitivity indices to those presented here. However, further work is required to develop methods to determine the number of simulator runs needed as well as suitable metrics to determine emulator quality. One aspect of this challenge is to develop optimal sizes for design data and test data so that emulators can be trained and evaluated in a way that minimises the number of simulator runs and optimises the emulator fit.

Emulators should be trained on design data that fill the input space evenly, and we chose to use Latin hypercube sampling in this study [40]. Other methods, such as orthogonal sampling [41] may provide a better sampling strategy.

The ability to evaluate emulators cheaply can be valuable for model calibration, where thorough exploration of a high dimensional input space is required [42]. A key benefit of a GP emulator approach is the explicit handling of uncertainty. Under the assumption that inputs and outputs have Gaussian distributions, the variance on the emulator output can be calculated directly, given variances on the inputs. This enables the direct calculation of sensitivity indices, as well as enabling a systematic investigation of the way that output uncertainties depend on uncertainties in the inputs.

### Limitations, challenges and future directions

The use of emulators to probe detailed biophysical models is at an early stage, and so there are several limitations and challenges associated with the present study.

#### Choice of inputs

We concentrated on the effect of maximum conductances in the present study, as this reduced the complexity of the analysis. The rationale for this approach was an assumption that kinetic parameters are determined by biophysics, and so less prone to variation than the expression of ion channels, pumps, and exchangers. However, a detailed sensitivity analysis of *I*_*Kr*_ dynamics in the Courtemanche model showed that these kinetic parameters influence APD [44], and other studies have highlighted difficulties in calibrating ion channel dynamics using traditional approaches as well as showing that different formulations can have an important effect on the magnitude and time course of an ion channel current [45]. In the present study our focus was on the action potential rather than *Ca*^2+^ handling, and a detailed sensitivity analysis of the mechanisms of *Ca*^2+^ storage, release, and uptake in each model would be a valuable extension to the work presented here. So far a fully comprehensive analysis has only been done for specially constructed models [15]. Nevertheless, a complete sensitivity analysis of biophysically detailed models, possibly using a hierarchical approach, remains an important challenge.

#### Simulator instability

One of the issues with a complete sensitivity analysis, highlighted in [15], is that parts of the simulator input space may generate implausible behaviours. For a cardiac cell model these behaviours might be a numerical instability, spontaneous beats, or failure to repolarise. In the present study we removed simulator runs from the design data where model behaviour was implausible, or where the simulator runs produced action potential alternans. We considered this to be a pragmatic approach. However, it is clearly an area for improvement because the location of these regions of input space conveys information about the model, and approaches where these locations are encoded explicitly show promise [46].

#### Choice of outputs

We selected a range of action potential features for our model outputs, these were based on measures used to describe experimental action potentials and aim to capture the main features of the action potential shape and duration. Our main focus was on the action potential. We included the maximum and minimum intracellular *Ca*^2+^ concentration as additional outputs, but did not consider the duration of the *Ca*^2+^ transient. We would not consider our choice of outputs to be definitive, and there may be better choices. A principal component analysis of the design data used in the present study showed that 95% of the output variance was accounted for by the first 6 principal components for the Courtemanche model and the first 5 for the Maleckar model. Parameterising the action potential and *Ca*^2+^ transient so that they are described by a minimal set of features is likely to be important not only for model analysis but also for model calibration [42]. Emulators that emulate time-dependent outputs have been developed, but are not yet widely used [47, 48], but could be a promising tool for extending work in this area.

#### Future directions

Extending the use of emulators from models of cardiac cells to models of cardiac tissue is an important next step, and initial studies are promising [49]. At present, tissue calculations are computationally expensive, especially for personalised meshes. However, the need to evaluate uncertainty in model predictions for use in the clinical setting requires computationally efficient approaches, and we anticipate exciting developments in this area.

## Supporting information

### S1 Text Mathematical details

Details of the procedures used to fit and evaluate the Gaussian process emulators, and to calculate main effects and sensitivity indices.

